# Benchmarking Gene Regulatory Network Inference Methods on Simulated and Experimental Data

**DOI:** 10.1101/2023.05.12.540581

**Authors:** Michael Saint-Antoine, Abhyudai Singh

## Abstract

Although the challenge of gene regulatory network inference has been studied for more than a decade, it is still unclear how well network inference methods work when applied to real data. Attempts to benchmark these methods on experimental data have yielded mixed results, in which sometimes even the best methods fail to outperform random guessing, and in other cases they perform reasonably well. So, one of the most valuable contributions one can currently make to the field of network inference is to benchmark methods on experimental data for which the true underlying network is already known, and report the results so that we can get a clearer picture of their efficacy. In this paper, we report results from the first, to our knowledge, benchmarking of network inference methods on single cell *E. coli* transcriptomic data. We report a moderate level of accuracy for the methods, better than random chance but still far from perfect. We also find that some methods that were quite strong and accurate on microarray and bulk RNA-seq data did not perform as well on the single cell data. Additionally, we benchmark a simple network inference method (Pearson correlation), on data generated through computer simulations in order to draw conclusions about general best practices in network inference studies. We predict that network inference would be more accurate using proteomic data rather than transcriptomic data, which could become relevant if highthroughput proteomic experimental methods are developed in the future. We also show through simulations that using a simplified model of gene expression that skips the mRNA step tends to substantially overestimate the accuracy of network inference methods, and advise against using this model for future *in silico* benchmarking studies.

## I. Introduction

### A. Network Inference Basics

In what is known as the “Central Dogma of Molecular Biology” [9], genetic information encoded in a cell’s DNA is transcribed into strands of messenger RNA (mRNA), which are then translated into proteins, which then carry out various functions within the cell. This process is also known as “gene expression.” Sometimes, a protein produced by one gene can positively or negatively affect the expression of another gene. For example, some proteins called transcription factors bind to the promoter regions of other genes and facilitate the attachment of RNA-polymerase, thereby having a positive effect on on the transcription of the gene. In this scenario, we say that the first gene “activates” the second gene. In other scenarios, a protein produced by one gene could block the promoter region of another gene, preventing its transcription. We refer to this interaction as “inhibition.” These positive and negative interactions between genes can form complex networks that play roles in signaling, response to stimuli, cell fate determination, and other important biological phenomena [43]. Disruptions to these networks can cause diseases, such as cancer [23].

Modern transcriptomic sequencing experiments such as single cell RNA-seq (scRNA-seq) [24], [39] allow us to to quantify the expression levels (in terms of mRNA abundance) of tens of thousands of genes, across hundreds or thousands of individual cells. A key research topic in computational biology is the attempt to leverage data from these mRNA sequencing experiments to infer the structure of the underlying gene regulatory networks producing them [22], [33]. This topic is known as “gene regulatory network inference”, or simply “network inference.” The strategy behind most network inference techniques is to calculate measures of statistical dependence, such as correlation or mutual information, between the mRNA levels of genes in a pair-wise fashion, and attempt to predict whether or not an interaction exists between the genes based on the strength of this measure. Many computational methods have been developed for this purpose [2], [3], [5], [7], [8], [14], [16], [19]–[21], [27], [28], [37], [38], [40]–[42]. Recently, network inference methods have been used to study a variety of problems in cell biology, including cancer [11]–[13], [30], [34], [35].

### B. Problem Formulation

We will now briefly give a formal definition of the network inference problem. Let us consider a situation in which we have *N* genes, and represent their expression levels with a set of random variables {*X*_1_, *X*_2_, …, *X*_*N*_ }. We represent each gene as a node in the network, and if a regulatory interaction exists (either activation or inhibition) between two genes *X*_*i*_ and *X*_*j*_, we represent it with the edge *X*_*i*_ → *X*_*j*_. For the purposes of this section, we do not differentiate between positive and negative edges. This is standard practice when it comes to evaluation methodology [26], as the difficult part of network inference is identifying whether a regulatory interaction exists not, and determining whether a known interaction is positive or negative is relatively easy.

We can then represent the network structure in matrix form. Let us consider a matrix *A*, with dimensions *N* × *N*. In this matrix there is a row for each gene, and a column for each gene. The element *A*_*i,j*_ (the *i*-th row and *j*-th column) is set to 1 if the edge *X*_*i*_→ *X*_*j*_ exists, and 0 if the edge does not exist.

So the matrix *A* represents the true network structure. The goal of network inference is to define a matrix *Â*, with dimensions *N* × *N*, in which each element *Â*_*i,j*_ represents our *prediction* about whether the edge *X*_*i*_ → *X*_*j*_ exists or not in the true network. Higher numbers mean that we are more confident that the edges exists, and lower numbers mean that we are less confident that it exists. Prediction values may be expressed as probabilities bounded on [0, 1], or they may not be.

Let us consider a quick example to illustrate these concepts. Figure 1 shows an example network with three genes, in which Gene 2 regulates Gene 1, and Gene 1 regulates Gene 3.

**Fig. 1.**
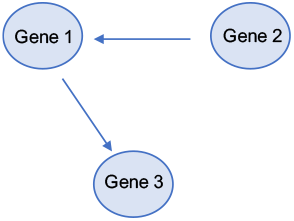
Example network with three genes, in which Gene 2 regulates Gene 1, and Gene 1 regulates Gene 3. Equation 1 shows the matrix representation of this network.

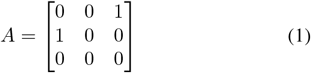

The above matrix *A* (Equation 1) represents the true network structure for the network shown in Figure 1. The element *A*_2,1_ = 1 tells us that that the edge *X*_2_→*X*_1_ exists, meaning that Gene 2 regulates Gene 1. The element *A*_1,2_ = 0 tells us that that the edge *X*_1_→ *X*_2_ does not exist, meaning that Gene 1 does not regulate Gene 2.

We then attempt to define a matrix *Â* with our network predictions. For example, we could define this matrix so that *Â*_*i,j*_= *f* (*X*_*i*_, *X*_*j*_) – this means that we calculate some function *f* of statistical dependence between each gene pair (*X*_*i*_, *X*_*j*_) where *i* ≠ *j*, and use its value as our prediction score. The matrix below in Equation 2 shows an example of what the network prediction matrix *Â* might look like.

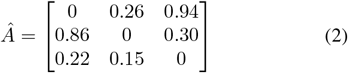

For the purposes of this paper, we are ignoring self-edges (*A*_*i,i*_). Although it is indeed possible for a gene to regulate itself [4], [29], uncovering these interactions requires more sophisticated experimental techniques and cannot be accomplished from statistical relationships between static measurements of gene expression. In this paper, we omit these self-edges and include only edges *X*_*i*_→ *X*_*j*_ where *i* ≠ *j* in the evaluation.

Some network inference methods, including methods based on Pearson correlation (which we will use throughout this paper), produce undirected network predictions. This means that in the network prediction matrix *A*_*i,j*_ = *A*_*j,i*_.

In other words, for every edge prediction *X*_*i*_ →*X*_*j*_, there is an equally weighted edge prediction *X*_*j*_→ *X*_*i*_, since the method does not try to predict the direction of the edge. In our benchmarking analysis, we will use a strict scoring protocol (more on this in the next subsection) in which an edge must be predicted in the correct direction to count as a true positive. This means that for every true positive scored by an undirected network prediction, it will also score a false positive (except in rare cases where two genes both regulate each other). With that being said, it is typically the case that the true underlying GRNs are quite sparse, with true edges making up only a small fraction of potential edges. So, if an undirected method is effective enough at identifying true edges, then it can still score well on the metrics described in the next subsection, even though most true positives scored will also score a false positive.

### C. Evaluation

Given an underlying true network structure matrix *A* and a network prediction matrix *Â*, generated with some network inference method, we can then evaluate the performance of the method by using two traditional machine learning metrics for binary classifier evaluation: receiver operating characteristic (ROC) curves and precision-recall (PR) curves.

Please recall that in the network prediction matrix *Â*, each element *Â* _*i,j*_ corresponds to our level of confidence that the edge *X*_*i*_→ *X*_*j*_ exists. We could apply a threshold to *Â* so that any elements above the threshold resolve to 1 and any elements below the threshold resolve to 0, giving us a concrete network prediction in which each edge is either predicted to exist or not. Higher thresholds applied to *Â* would give us more conservative predictions, as only the higher-confidence predicted edges would be kept in the network prediction. Conversely, a lower threshold would give us a less conservative prediction.

For any given threshold, we can compare the final predicted network to the true network *A*, and fill in the following table with the number of true positives, false positives, false negatives, and true negatives:

From this table, we can write the true positive rate (TPR) and false positive rate (FPR) as follows:

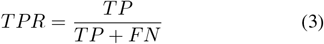

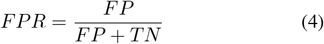

We want to simultaneously have a high true positive rate and a low false positive rate. However, this involves a tradeoff. Setting a low threshold helps us to increase our true positive rate, but also increases the false positive rate. Setting a high threshold helps us to decrease the false positive rate but also decreases the true positive rate. So, the choice of a threshold is somewhat arbitrary and depends on the researcher’s subjective preference about how to navigate the trade-off between true positives and false positives.

The key advantage of receiver operating characteristic (ROC) curves is that they give us a measure of the accuracy of the network inference method, considering all possible thresholds that could be applied to the prediction matrix *Â*. To plot a ROC curve, we simply plot the true positive rate against the false positive rate, across all of the possible thresholds. The area under this curve (called AUROC for short) gives us a single numerical score of the method’s accuracy, taking into account all possible thresholds. A perfect AUROC score is 1. Figure 2 shows an example of a ROC curve.

**Fig. 2.**
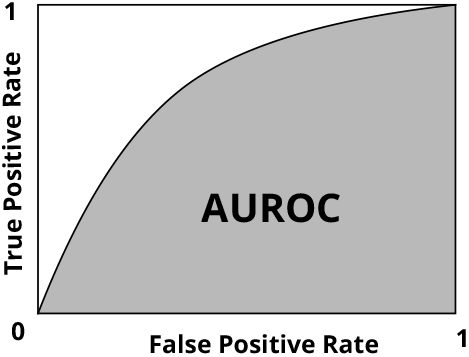
Example of a receiver operating characteristic (ROC) curve. The true positive rate is plotted against the false positive rate for every possible edge-weight threshold. The area under the curve (called AUROC) gives a numerical measure of accuracy. A perfect AUROC score is 1.

In order to have a point of comparison, we can define a “no skill control” ROC curve, using a classifier that simply guesses the most common label for every item. In the case of network inference, this means guessing that every possible edge does not exist. In other words, we define a prediction matrix *Â*, for which *Â* _*i,j*_ = 0 for every value of *i* and *j* such that *i* ≠ *j*. The AUROC of the no skill control ROC curve is always 0.5, by definition. Notably, this is also the AUROC one would get, on average, by randomly guessing edge weights so that all potential edge predictions are randomly ordered in terms of the prediction confidence. So, if a network inference method achieves an AUROC score higher than 0.5, we can say that it has outperformed the no skill control, or that it has outperformed the average AUROC that would be expected by random guessing.

We can also use a similar measure called a precision-recall (PR) curve to score accuracy. To do this, we define precision and recall as follows, again using the numbers from Table I:

**TABLE I.** Evaluation Table

|  | Edge Exists | Edge Does NOT Exist |
| --- | --- | --- |
| Edge Predicted | True Positive (TP) | False Positive (FP) |
| Edge NOT Predicted | False Negative (FN) | True Negative (TN) |

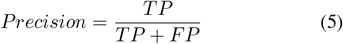

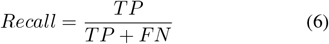

You may notice that recall is the same measure as the true positive rate, and is just given a different name, somewhat confusingly, when plotted on a precision-recall curve. To plot the curve, we plot the precision against the recall, across all possible thresholds. As with the ROC curve, we can also use the area under the PR curve (called AUPR for short) as a single numerical measure of accuracy, for which 1 is a perfect score. Figure 3 shows an example of a PR curve.

**Fig. 3.**
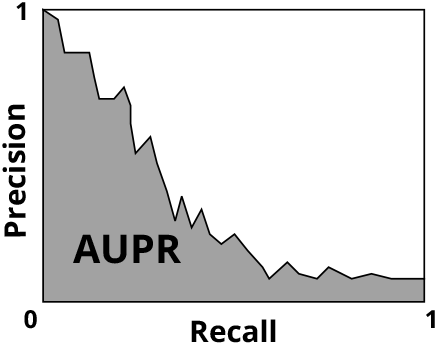
Example of a precision-recall (PR) curve. The precision is plotted against the recall (same as the true positive rate) for every possible edgeweight threshold. The area under the curve (called AUPR) gives a numerical measure of accuracy. A perfect AUPR score is 1.

We can define a “no skill control” AUPR to serve as a point of comparison. This no skill control AUPR is simply the number of edges that actually exist in the true network divided by the number of possible edges. this is also the AUPR one would get, on average, by randomly guessing edge weights so that all potential edge predictions are randomly ordered in terms of the prediction confidence. So if a network inference method achieves a higher AUPR than the no skill control AUPR, we can say that it has outperformed the average AUPR that one would expect from random guessing. While the no skill control AUROC is 0.5 by definition, different networks can have different no skill control AUPR scores, depending on how many true edges they have and how many possible edges they have.

Network inference methods can be evaluated by testing them on data for which the structure of the underlying network is already known, and seeing how well they can predict the true network structure. Often the data used for evaluation comes from a model organism, such as *E. coli* or *S. cerevisiae* (yeast). Previous benchmarking attempts on model organisms have yielded mixed results. For example in the famous 2012 paper “Wisdom of Crowds for Robust Gene Network Inference” [26], the network inference methods tested performed reasonable well on the *E. coli*, but failed to considerably outperform random guessing on the *S. cerevisiae* data. A similar mixed result was found in [32]. Recent theoretical work has explored how network inference from transcriptomic data may be more or less feasible under different conditions. Mahajan et al. 2022 [25] shows through simulations and mathematical analysis that, under conditions of only intrinsic noise in the processes of transcription and translation, the correlation between mRNA abundance and protein abundance *even for the same gene* becomes quite weak if there is a large difference between the mRNA stability and protein stability. [25] also shows that under these conditions the mRNA abundance for transcription factor and target genes in a regulatory relationship become uncorrelated when there is a large difference between mRNA and protein stability. This is a pessimistic result for the challenge of network inference, since most methods rely on the assumption that mRNA abundance can be used as a reliable proxy for protein abundance, and that regulatory interactions between genes can be detected based on statistical dependencies between mRNA levels. However, as we noted previously, [25] assumes only intrinsic noise in gene expression. New results in [31] show that the network inference problem may be more tractable under conditions of extrinsic noise in the gene regulatory relationship.

It is also worth noting that many of the previous benchmarking studies, including [26] and [32], used microarray expression data, not single cell RNA-seq data, and it is not clear if methods that perform well on one will necessarily perform well on the other. So, one of the most pressing problems in the study of network inference is to simply continue benchmarking methods on real experimental data to improve our understanding of how well they perform under different conditions, in different organisms, and using different experimental methods.

Ultimately, what matters is how well network inference methods perform on real biological datasets, so benchmarking on data from model organisms is preferable to benchmarking on data from simulations, which may contain the researchers’ own assumptions about the biological networks, leading to an overestimate of the method’s accuracy (more on this later). With that being said, *in silico* benchmarking can still be useful, as it allows researchers to investigate how network inference methods perform on simulations of specific network motifs, parameter sets, noise conditions, etc. In this paper we will use both types of benchmarking to attempt to answer different questions. We will use *in silico* data from simulations to answer questions about best practices for network inference studies, including the hypothetical question of how well network inference on proteomic data would work compared to transcriptomic data, and the question of whether or not it is advisable to use a simplified one-step model of gene expression in *in silico* benchmarking studies. We will use experimental single cell *E. coli* data to test whether network inference methods can uncover known gene regulatory relationships with more accuracy than random guessing, and whether published methods known to perform well on microarray data also perform well on single cell data.

## II. In-Silico Benchmarking

In this section, we report results of an *in silico* benchmarking analysis. The goal of this section is primarily to use results from simulations to draw conclusions about best practices in network inference research. Consistent with previous work in [25] and [31], we use Pearson correlation (a linear measure) to quantify the statistical dependence between mRNA abundance levels. We also performed this analysis with two nonlinear measures (Spearman rank correlation and mutual information), but those results are omitted from this paper because they were so similar to the Pearson correlation results as to not be especially interesting.

We report two main results related to best practices in network inference research. The first result is that one can expect a gain in accuracy by performing network inference on proteomic, rather than transcriptomic, data. Currently, network inference analyses are almost always performed on transcriptomic data, because modern high-throughput experimental techniques can more easily measure mRNA abundance than protein abundance. However, in the future it may be feasible to collect protein abundance measurements in a high-throughput manner [10].

The second result reported in this section is related to best practices for *in silico* benchmarking of network inference methods. The process of gene expression involves two steps: DNA → mRNA → Protein. However, sometimes in *in silico* benchmarking research a simplified model of gene expression is used, in which the intermediate step is ommitted: DNA → Gene Product. The “Gene Product” is then used to represent both mRNA and protein abundance. We show, through simulations, that using this type of simplified model of expression for benchmarking tends to overestimate the accuracy, and therefore is not advisable for *in silico* benchmarking efforts.

### A. Simulating a Network

We generated our *in silico* datasets by running stochastic simulations of a network with 20 genes. The network structure is shown in Figure 4. The first 10 genes, called G1, G2, …, G10, are connected in a topology that we took from a subnetwork of the *E. coli* GRN reported by Fang et al. 2017 [15], so as to have a realistic topology. The other 10 genes, called G11, G12, …, G20, are decoys that are not connected to any other genes.

**Fig. 4.**
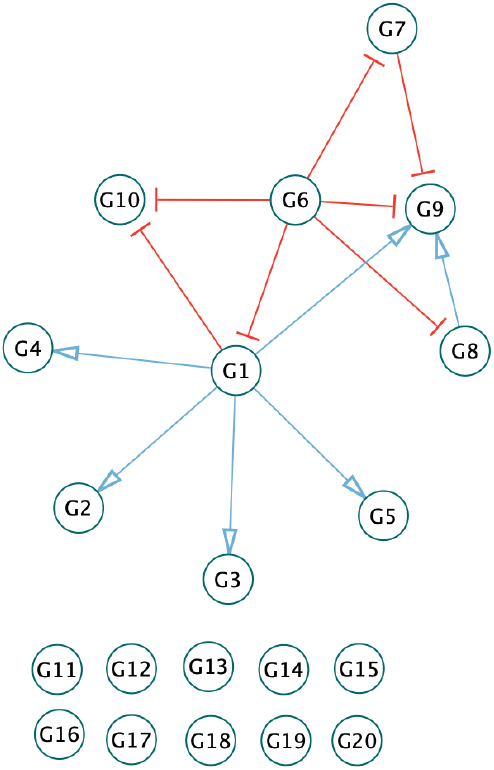
Visual representation of the network of 20 genes we used for our *in silico* simulations. Blue lines show activation (positive) regulatory relationships, while red lines show inhibition (negative) regulatory relationships. The first 10 genes (G1, G2,…,G10) are connected in a topology that we took from a subnetwork of the *E. coli* GRN reported by Fang et al. 2017 [15]. The other 10 genes (G11, G12, …, G20) are decoys that do not have any regulatory interactions with any other genes.

For each gene, we modeled the production and degradation of both the mRNA and the protein, and we modeled the regulatory interactions between genes with Hill functions [1]. We will not list the entire mathematical model here (which involves 40 variables, tracking the mRNA and protein counts for the 20 genes). Rather, we will give brief examples to highlight the key concepts of the model.

First, we consider the case of a gene that is not regulated by any other gene. Table II shows the stochastic model we can use to track the mRNA count (*M*) and protein count (*P*) for this situation, where a gene is being expressed with no regulation. In this model, *k* is the mRNA production rate, *γ* is the mRNA degradation rate, *k*_*p*_ is the protein production rate, and *γ*_*p*_ is the protein degradation rate.

**TABLE II.** No Regulation Model

| Event | Count Update | Propensity |
| --- | --- | --- |
| mRNA1 production | $M \rightarrow M + 1$ | $k$ |
| mRNA1 degradation | $M \rightarrow M - 1$ | $\gamma M$ |
| Protein production | $P \rightarrow P + 1$ | $k_p M$ |
| Protein degradation | $P \rightarrow P - 1$ | $\gamma_p P$ |

Let us now consider a scenario in which the expression of one gene, which we will call Gene B, is activated by another gene, which we will call Gene A. Table III shows the stochastic model that we can use to track the counts of the Gene A mRNA (*M*_*a*_), Gene A protein (*P*_*a*_), Gene B mRNA (*M*_*b*_) and Gene B protein (*P*_*b*_). It is similar to the model described in Table II, except that it includes two genes, and the rate of transcription for the second gene is given by the Hill function 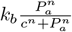, which depends on the protein count of the first gene, and uses the constant parameters *c* and *n*.

**TABLE III.** Activation

| Event | Count Update | Propensity |
| --- | --- | --- |
| mRNA A production | $M_a \rightarrow M_a + 1$ | $k_a$ |
| mRNA A degradation | $M_a \rightarrow M_a - 1$ | $\gamma_a M_a$ |
| Protein A production | $P_a \rightarrow P_a + 1$ | $k_{pa} M_a$ |
| Protein A degradation | $P_a \rightarrow P_a - 1$ | $\gamma_{pa} P_a$ |
| mRNA B production | $M_b \rightarrow M_b + 1$ | $k_b \frac{P_a^n}{c^n + P_a^n}$ |
| mRNA B degradation | $M_b \rightarrow M_b - 1$ | $\gamma_b M_b$ |
| Protein B production | $P_b \rightarrow P_b + 1$ | $k_{pb} M_b$ |
| Protein B degradation | $P_b \rightarrow P_b - 1$ | $\gamma_{pb} P_b$ |

In order to model a scenario in which Gene A is inhibiting, rather than activating Gene B, we use the model described in Table III, except that we change the rate of mRNA B production from 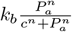 to 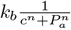, to model the negative regulation. For scenarios in which a gene is regulated by more than one other gene, we make the simplifying assumption that the regulatory effects can be modeled separately and summed together.

To simulate the experimental workflow of a gene expression measurement experiment, we ran 100 simulations of the network using Gillespie’s stochastic simulation algorithm [18], with each run representing an experimental sample, and recorded the counts of mRNA and protein for each gene, for each run. We then performed the network inference, calculating the pair-wise Pearson correlation scores for each pair of genes. We performed this Pearson correlation network inference once for the mRNA data and once for the protein data. We note that Pearson correlation is a linear correlation measure, and one of the simplest network inference methods. We also performed this analysis with two nonlinear measures (Spearman rank correlation and mutual information), but those results are omitted from this paper because they were so similar to the Pearson correlation results as to not be especially interesting.

We then calculated the AUROC and AUPR accuracy scores, described in the previous section, for both the mRNA network inference prediction and protein network inference prediction. We repeated this process 20 times to generate distributions of the AUROC and AUPR scores, which were used to calculate the error bars in the figures. Lastly, we repeated this entire analysis for different assumptions about the relative protein stability compared to mRNA stability. Recent theoretical findings from [25] suggest that the challenge of network inference is more difficult when there is a large difference between protein and mRNA stability, so this stability ratio is relevant for our *in silico* benchmarking study.

### B. Network Inference from mRNA and Protein Measurements

The first topic we will address is the difference between the accuracy of network inference from transcriptomic (mRNA abundance) and proteomic (protein abundance) datasets. In practice, gene regulatory network inference is almost always performed using transcriptomic data from experiments such as RNA-seq. However, in the future it may be feasible to collect protein abundance measurements in a high-throughput manner [10]. Would the development of these proteomic experimental methods be useful from a network inference standpoint?

Figure 5 shows the AUROC scores of network predictions generated using Pearson correlation, for different protein/mRNA stability ratios (please note that the horizontal axis is in base-2 log scale). The red dots show results for network inference using protein abundance data, and the blue dots show the results for mRNA abundance data. Error bars for both show one standard deviation. Figure 6 is a similar plot, but with AUPR scores rather than AUROC scores. Again, red dots show results for network inference using protein data, blue dots show results for mRNA data, and error bars show one standard deviation.

**Fig. 5.**
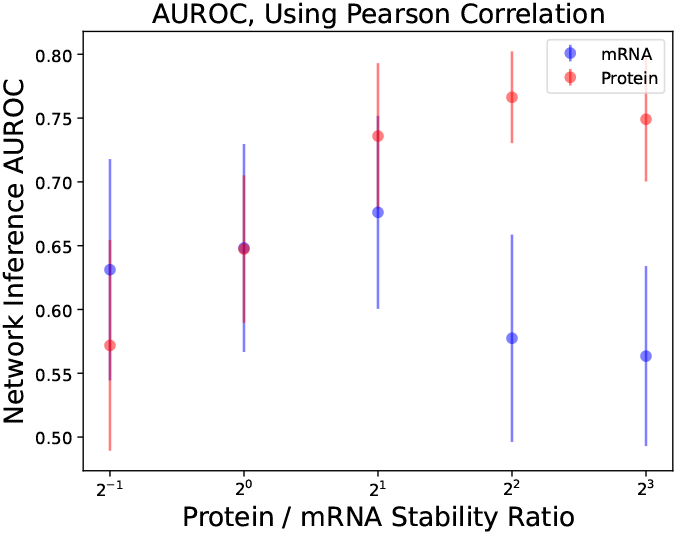
Area under the receiver operating characteristic curve (AUROC) scores for network inference predictions generated with Pearson correlation. Blue dots show results using mRNA abundance data, and red dots show results using protein abundance data. Error bars show one standard deviation.

**Fig. 6.**
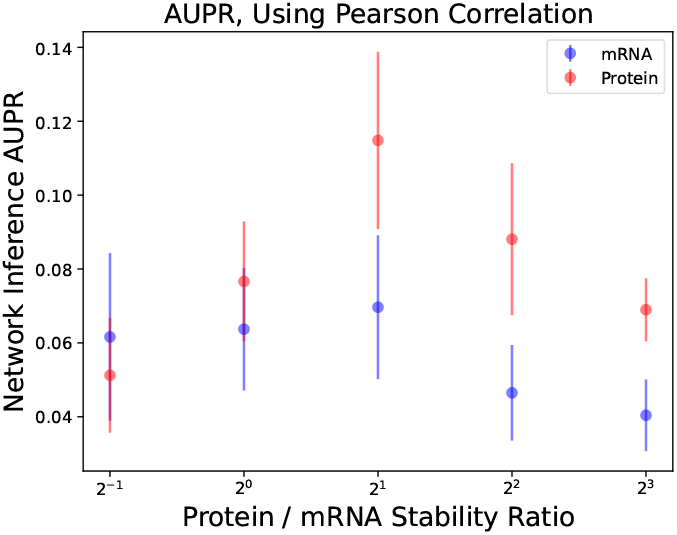
Area under the precision-recall curve (AUPR) scores for network inference predictions generated with Pearson correlation. Blue dots show results using mRNA abundance data, and red dots show results using protein abundance data. Error bars show one standard deviation.

We note that in both Figure 5 and Figure 6, network inference using the more stable molecule type tended to yield greater accuracy. This difference in accuracy is especially pronounced for the protein/mRNA stability ratios of 2^1^, 2^2^, and 2^3^, suggesting that using proteomic data could yield substantial gains in accuracy over transcriptomic data, even when the proteins are only moderately more stable relative to the mRNAs.

Although it is less common biologically, we also simulated a scenario in which the mRNAs were more stable relative to the proteins (stability ratio of 2^−1^ on the horizontal axis). In this scenario, network inference from mRNA data yielded slightly more accurate results on average than network inference from protein data, although there was substantial overlap between the two.

The key insight here is that, in the context of network inference from real-world biological datasets, one can probably expect more accurate results using protein abundance data rather than mRNA abundance data, and it is better to use the protein data if it is available. This is assuming that the proteins are relatively more stable than the mRNA, which is typically the case [1]. Currently, mRNA data is much more readily available than proteomic data, but if high-throughput proteomic measurement techniques are developed in the future, then that could considerably increase the accuracy and feasibility of gene regulatory network inference.

### C. Omitting mRNA Step Over-estimates Method Accuracy

The next topic we sought to investigate through simulation was the use of a simplified model of gene expression for *in silico* network inference benchmarking. In the previous subsection, we were running simulations using a stochastic model that included both the mRNA and protein steps of gene expression: DNA → mRNA →Protein. However, sometimes in *in silico* benchmarking research a simplified model of gene expression is used, in which the intermediate step is ommitted: DNA→ Gene Product. In the simplified model, only the production and degradation of the Gene Product is simulated, and the Gene Product level is used as a proxy for both the mRNA and the protein, depending on the situation.

It stands to reason that this simplified model of gene expression is likely to overestimate the accuracy of network inference methods when used for *in silico* benchmarking. However, to our knowledge, this has not previously been formally demonstrated and quantified. In this section, we introduce a simplified version of the stochastic simulation model described previously, in which the mRNA step is omitted. In this model, Table II is rewritten as Table IV. Rather than tracking the mRNA counts (*M*) and protein counts (*P*) separately, we instead use the stochastic process *G* to track the count of the “Gene Product”, which is purposely defined ambiguously so that it can refer to either the mRNA or the protein.

**TABLE IV.** No Regulation Simplified Model

| Event | Count Update | Propensity |
| --- | --- | --- |
| Gene product production | $G \rightarrow G + 1$ | $k$ |
| Gene product degradation | $G \rightarrow G - 1$ | $\gamma G$ |

For the scenario in which one gene (Gene A) activates another (Gene B), Table III is rewritten as Table V.

**TABLE V.** Activation

| Event | Count Update | Propensity |
| --- | --- | --- |
| Gene A product production | $G_a \rightarrow G_a + 1$ | $k_a$ |
| Gene A product degradation | $G_a \rightarrow G_a - 1$ | $\gamma_a G_a$ |
| mRNA B production | $G_b \rightarrow G_b + 1$ | $k_b \frac{G_a^n}{c^n + G_a^n}$ |
| mRNA B degradation | $G_b \rightarrow G_b - 1$ | $\gamma_b G_b$ |

Figure 7 shows the AUROC scores of network predictions generated using Pearson correlation, with simulation data generated from the simplified model of gene expression omitting the mRNA step. These results are shown alongside the previous results from Figure 5, to serve as a point of comparison. Note that the AUROC scores for the simplified model were higher, on average, than both the mRNA data and protein data AUROC scores from the previous section for every protein / mRNA stability ratio (although in some cases there was considerable overlap with the AUROC scores from protein data). Figure 8 shows a similar result (including the previous results from Figure 6), except with AUPR scores instead of AUROC scores. Again, the AUPR scores for the simplified model were higher, on average, than than the scores for the more realistic model of gene expression (although again, there was some overlap with the scores from proteomic data), for every protein/mRNA stability ratio.

**Fig. 7.**
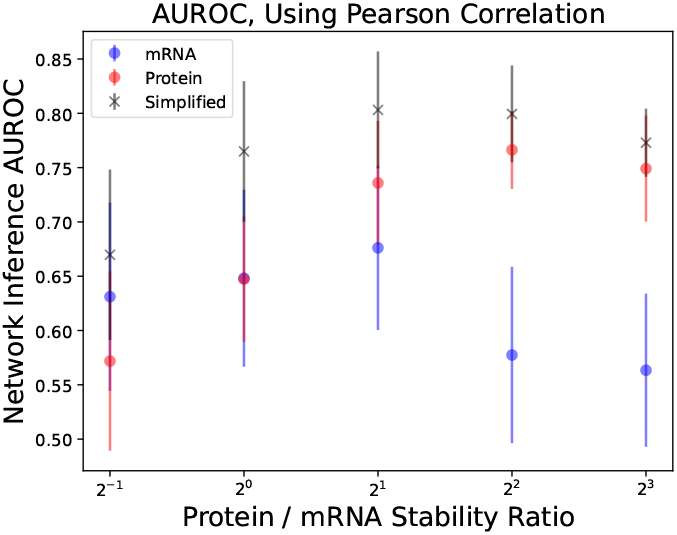
Area under the receiver operating characteristic curve (AUROC) scores for network inference predictions generated with Pearson correlation. Blue dots show results using mRNA abundance data, and red dots show results using protein abundance data. X-marks show results for simplified model, omitting the mRNA step. Error bars show one standard deviation.

**Fig. 8.**
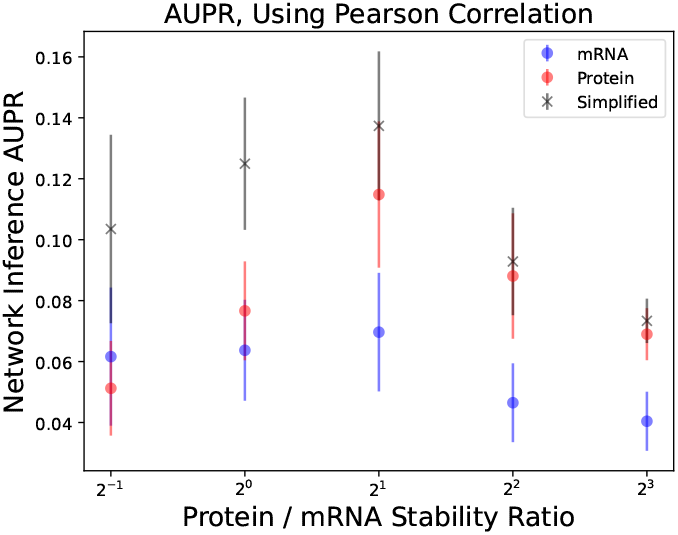
Area under the precision-recall curve (AUPR) scores for network inference predictions generated with Pearson correlation. Blue dots show results using mRNA abundance data, and red dots show results using protein abundance data. X-marks show results for simplified model, omitting the mRNA step. Error bars show one standard deviation.

These results suggest that using the simplified model without the mRNA step will tend to overestimate the accuracy of network inference methods compared to a more realistic model of gene expression (even assuming a best-case scenario in which proteomic data is available), and so the simplified model of gene expression generally should not be used for benchmarking and method validation if it is feasible to use the more realistic model instead. Ultimately, the performance of network inference methods can only truly be validated on real experimental datasets (as we will do in the next section), but if experimental data is unavailable and it is necessary to generate data through simulations, then it is important to use a realistic model of gene expression that includes both the transcription and translation steps to avoid overestimating the method’s accuracy.

## III. Experimental Data Benchmarking

In the previous section, we used simulations of a gene regulatory network to draw conclusions about best practices for network inference studies, including the advantages of using proteomic data rather than mRNA data, and problems that can result from using a simplified one-step model of gene expression for *in silico* benchmarking. Although working with simulated *in silico* data can be useful and yield valuable insights, what ultimately matters the most is the performance of network inference techniques on real biological systems. In this section, we benchmark network inference techniques on single cell mRNA abundance data from *E. coli* [6], in an attempt to address two questions.

The first question is whether it is possible to predict gene regulatory interactions with accuracy considerably greater than random guessing, based on statistical dependencies mRNA levels for this cell type (*E. coli*) and experimental protocol (single cell PETRI-seq [6]). Although this may seem like a low bar, success on this task constitutes a nontrivial result since, as we noted earlier, previous benchmarking attempts [26], [32] have yielded mixed results in which sometimes even the best methods have failed to considerably outperform random guessing, while other times performing quite well. Furthermore, previous theoretical work suggests that there are some conditions in which network inference from mRNA data may be possible and other conditions in which it may be impossible [25], [31].

The second question we hope to address is related to the performance of published network inference methods that were previously validated on microarray data. Most previous benchmarking efforts have used microarray data [26], [32], and we are interested to see if impressive accuracy on microarray data translates to impressive accuracy on single cell data. We benchmark GENIE3 [20], which was regarded as one of the best network inference methods for use on microarray data, as well as CLR [14], an older method published in 2007. To serve as a point of comparison, we also include PIDC [8], a relatively new method (published in 2017), developed for application on single cell data. We also include three simplistic methods: Pearson correlation, mutual information, and Spearman rank correlation, with the idea being that for a published method to show truly impressive accuracy on this dataset, it must not only outperform the no skill control, but must also outperform these simplistic methods that can be easily implemented with a few lines of code.

### A. Experimental Dataset

In this section, we benchmark network inference techniques on single cell mRNA abundance data from *E. coli*, using data from Blattman et al. 2019 [6]. To our knowledge, this is the first benchmarking of network inference methods on single cell *E. coli* (previous benchmarking was done on microarray and bulk RNA-seq data). This is because until recently single-cell RNA-seq methods were not amenable to prokaryotic cells, due to their low mRNA copy numbers, lack of mRNA polyadenylation, and thick cell walls [6]. However, Blattman et al. developed a clever new experimental method for single cell RNA sequencing in prokaryotes, using in situ combinatorial indexing to barcode transcripts in tens of thousands of cells before sequencing. They have named this method PETRI-seq (Prokaryotic Expression profiling by Tagging RNA In situ and SEQuencing).

We downloaded data from three of the Blattman et al. 2019 [6] *E. coli* sequencing experiments, called Experiment 1.06, Experiment 1.10, and Experiment 1.12, all available on the Gene Expression Omnibus under accession number GSE141018. The available data was in the form of raw counts, so we performed TPM normalizations for the three datasets. We then performed an additional data processing step, removing any genes that did not have any recorded expression, and any samples that did not have any recorded expression. It is interesting to note that, unlike the microarray datasets used in previous benchmarking efforts [26], [32], the PETRI-seq experimental datasets are quite sparse, with many genes showing zero recorded expression for many cells, and relatively uncommon occurrences of recorded expression. This difference may be problematic for network inference methods developed for use on microarray data.

For our “ground-truth” *E. coli* gene regulatory network, we used the GRN reported by Fang et al. 2017 [15]. This is referred to as the high confidence transcriptional regulatory network (hiTRN) in their paper, and includes gene regulatory interactions from RegulonDB [17], as well as several additional interactions supported by ChIP-seq evidence.

For our final data processing step, we selected the genes in the experimental datasets that overlapped with the set of genes in the gold-standard network (thereby excluding any genes in the experimental datasets that did not have at least one interaction according to the gold standard network). Descriptions of these datasets after processing are shown in Table VI.

**TABLE VI.**
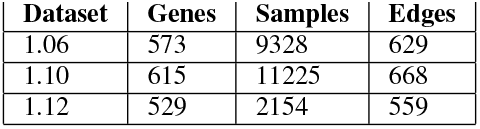
Dataset Descriptions

| Dataset | Genes | Samples | Edges |
| --- | --- | --- | --- |
| 1.06 | 573 | 9328 | 629 |
| 1.10 | 615 | 11225 | 668 |
| 1.12 | 529 | 2154 | 559 |

### B. Network Inference Methods

We then used the three datasets described in the previous subsection to infer the underlying network using six network inference methods: Pearson correlation, mutual information, Spearman correlation, GENIE3 [20], PIDC [8], and CLR [14].

The first three methods are simple approaches that can be easily implemented in any programming language, and do not involve any special software. Pearson correlation is the simplest method, a linear measure of correlation, and is useful as a point of comparison to see if the more complex and elaborate methods outperform it in terms of accuracy. Mutual information is a non-linear measure of statistical dependence based on information theory [36]. Spearman correlation is a rank-based nonlinear correlation measure.

The next three methods are examples of more complex, published network inference methods. GENIE3 [20] is a network inference method that uses a tree-based regression technique. It was published in 2010 and was considered to be one of the strongest and most accurate network inference methods at the time. However, it predates the popularization of single cell RNA-seq experiments and was developed primarily for use on microarray gene expression datasets, so it was not clear how well it would perform on single cell data. PIDC [8] is a network inference method based on information theory. It is similar to mutual information, except that the information-theoretic dependencies are computed in a triplet-wise rather than pair-wise manner. PIDC is relative new (published in 2017), and was developed for use on single cell data. The context likelihood of relatedness (CLR) algorithm [14], originally published in 2007 and validated on microarray data, is another information-theoretic approach. Here, we use the CLR implementation available in the PIDC code package [8].

### C. Results

We scored the network predictions for each method and each replicate against the true underlying network using the receiver operating characteristic (ROC) and precision-recall (PR) curve methods described earlier in this paper. Figure 9 shows the resulting ROC curves for each experimental replicate, using Pearson correlation, mutual information, and Spearman correlation. A line showing the no skill control ROC curve is also included. Note that the network inference ROC curves modestly outperform the no skill control curve, although they are still far from perfect.

**Fig. 9.**
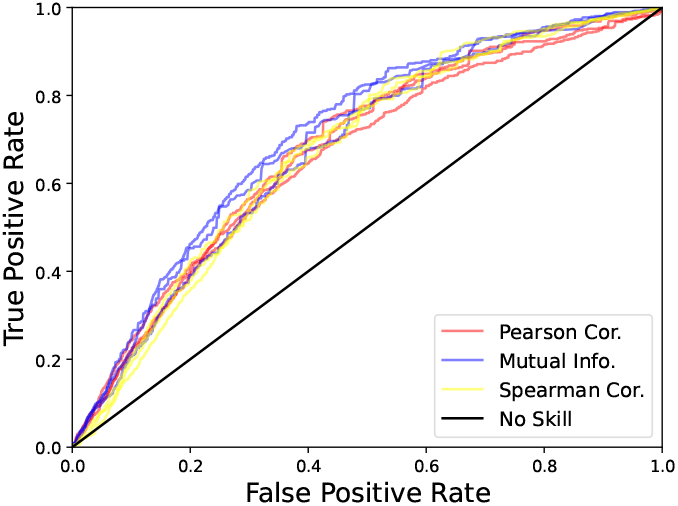
Receiver operating characteristic (ROC) curves for Pearson correlation (red), mutual information (blue), Spearman correlation (yellow), and the no skill control (black). Curves are shown for each method and for each of the three single cell PETRI-seq *E. coli* replicates [6] (experiments 1.06, 1.10, and 1.12). All of the network inference methods outperform the no skill control and show about the same level of accuracy, with mutual information being very slightly better than Pearson correlation and Spearman correlation.

Figure 10 shows the resulting ROC curves for each replicate, for for Pearson correlation, GENIE3 [20], PIDC [8], and CLR [14], as well as the no skill control line (shown in black). This is a very similar plot to Figure 9, except that the methods have been split up so as to not crowd the figure with too many curves. However, the Pearson correlation ROC curves have been kept in Figure 10 to serve as a point of comparison. Note that the curves for Pearson correlation, PIDC, and CLR appear to outperform the no skill control for all of the replicates, while for GENIE3 it is less clear.

**Fig. 10.**
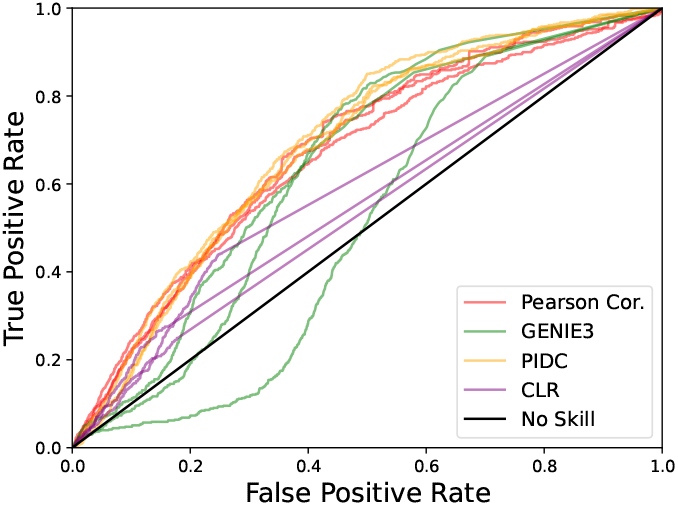
Receiver operating characteristic (ROC) curves for Pearson correlation (red), GENIE3 [20] (green), PIDC [8](orange), CLR [14] (purple), and the no skill control (black). Curves are shown for each method and for each of the three single cell PETRI-seq *E. coli* replicates [6] (experiments 1.06, 1.10, and 1.12). The curves for Pearson correlation, PIDC, and CLR appear to outperform the no skill control for all of the replicates, while for GENIE3 it is less clear.

Figure 11 shows the mean area under the ROC curve (AUROC) for each method, calculated across the three datasets, with error bars showing one standard deviation. We have also included a bar showing the area under the no skill control ROC curve, which is 0.5 by definition. Figure 12 shows a similar bar plot, but with the mean area under the precision-recall curves (AUPR), calculated across the three datasets, with error bars showing one standard deviation. Table VII and Table VIII show the AUROC and AUPR scores, respectively, for each of the methods and each of the replicates. Note that the network inference method results are very far from perfect, but considerably better than one would expect from random guessing (represented by the no skill scores). For both the AUROC scores and the AUPR scores, all of the network inference methods outperformed the no skill control, but GENIE3 [20] and CLR [14] did not perform as well as the more simplistic methods. Mutual information performed the best overall, for both the AUROC and the AUPR scores.

**TABLE VII.**
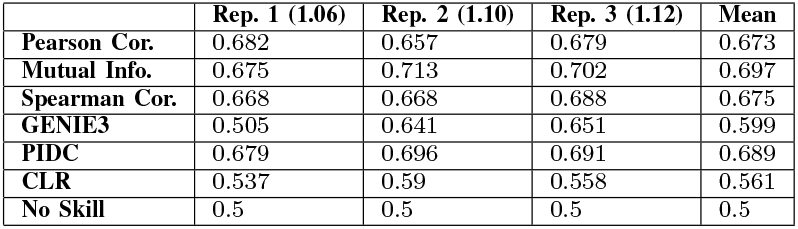
AUROC Table

**TABLE VIII.** AUPR Table

|  | Rep. 1 (1.06) | Rep. 2 (1.10) | Rep. 3 (1.12) | Mean |
| --- | --- | --- | --- | --- |
| **Pearson Cor.** | 0.0038 | 0.003 | 0.0036 | 0.0035 |
| **Mutual Info.** | 0.0038 | 0.0037 | 0.004 | 0.0038 |
| **Spearman Cor.** | 0.0032 | 0.0028 | 0.0035 | 0.0032 |
| **GENIE3** | 0.0019 | 0.0024 | 0.003 | 0.0024 |
| **PIDC** | 0.0033 | 0.003 | 0.0036 | 0.0033 |
| **CLR** | 0.0025 | 0.0025 | 0.0029 | 0.0026 |
| **No Skill** | 0.0019 | 0.0018 | 0.002 | 0.0019 |

**Fig. 11.**
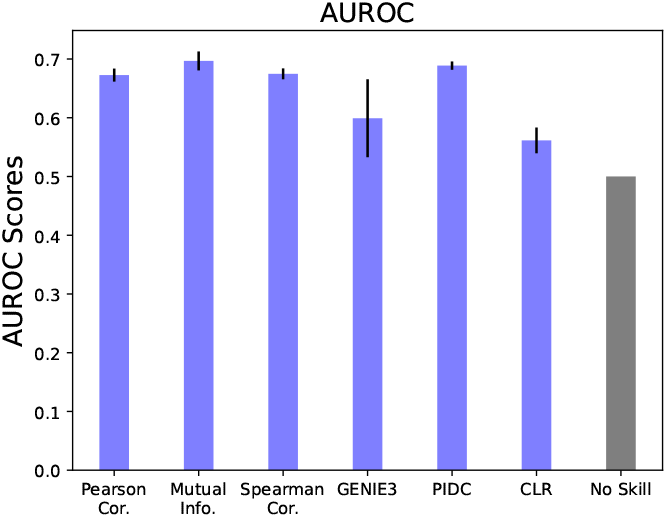
Bars show mean area under the reciever operating characteristic curve (AUROC) scores for each method, computed across the three single cell PETRI-seq *E. coli* replicates [6] (experiments 1.06, 1.10, and 1.12). Error bars show one standard deviation. A bar for the no skill control AUROC (which is 0.5 by definition) is also shown for comparison. All of the network inference methods outperformed the no skill control, but GENIE3 [20] and CLR [14] did not perform as well as the more simplistic methods.

**Fig. 12.**
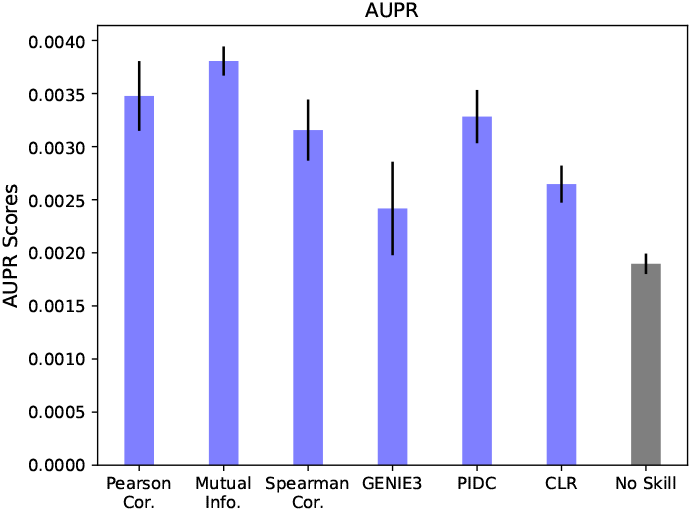
Bars show mean area under the precision-recall curve (AUPR) scores for each method as well as the no skill control, computed across the three single cell PETRI-seq *E. coli* replicates [6] (experiments 1.06, 1.10, and 1.12). Error bars show one standard deviation. All of the network inference methods outperformed the no skill control, but GENIE3 [20] and CLR [14] did not perform as well as the more simplistic methods. Mutual information had the best performance.

## IV. Discussion

In this paper, we have presented network inference benchmarking results using both simulated and experimental datasets. A key conclusion from the simulation benchmarking is that one can expect substantial gains in network inference accuracy by using proteomic rather than transcriptomic datasets (assuming that the proteins are more stable relative to the mRNAs – in rare cases where the mRNAs are more stable, transcriptomic data should be used). Currently it is difficult to find proteomic datasets that are usable for network inference, but this may change in the future as highthroughput proteomic experimental methods are developed [10]. Another conclusion from the simulation benchmarking is that it is not advisable to use the simplified model of gene expression, in which the mRNA step is omitted, for the purpose of benchmarking network inference methods as making this simplification tends to overestimate the accuracy of the network inference method being tested.

In addition to the simulation data benchmarking, this paper also contains, to our knowledge, the first benchmarking of network inference methods on single cell *E. coli* transcriptomic data (since it was only recently that an RNA-seq experimental protocol suitable for use on prokaryotes was developed). Although the topic of network inference has been widely studied for more than a decade, attempts to validate methods on real datasets have been insufficient, and have produced mixed results in which methods sometimes work well and sometimes fail to outperform random chance. So, one of the most valuable contributions one can currently make to the field of network inference is to simply test out existing methods on real experimental datasets and report the results, so that we can get a clearer picture of how effective these methods are.

The results of our benchmarking on experimental data can be interpreted differently based on how high we set the bar for effectiveness. If a network inference method must reliably and consistently be able to uncover the true network structure in order to be considered effective, then the methods tested in this paper fail to meet this standard. Indeed, even when constructing a network prediction with only the highest-confidence predicted edges, one may still end up with more false positives than true positives. However, if we define effectiveness as the ability of a method to consistently and reliably outperform random chance, then the methods tested in this paper *do* meet this standard. Although far from perfect, this level of accuracy may still be useful for biologists in terms of hypothesis generation. For example, if an experimental biological is planning a knockout experiment, they may find network inference to be a valuable tool for choosing targets and generating hypotheses about regulatory interactions, which can then be tested experimentally.

Another key result from our benchmarking on experimental data is that published methods may perform differently on single cell datasets compared to bulk and microarray datasets. For example, GENIE3 was considered to be one of the strongest and most accurate methods for microarray data, but failed to outperform the much simpler methods of Pearson correlation, mutual information, and Spearman correlation on the single cell *E. coli* data. This result is notable because many of the most widely used network inference methods were initially validated on microarray or bulk RNA-seq data, and may need to be re-evaluated if they are to be used on single cell datasets.

This paper is not meant to be a definitive, conclusive statement on the efficacy of network inference from transriptomic data. On the contrary, we believe that due to the mixed results reported by previous studies, one of the most important research goals for the network inference topic is the continued benchmarking of methods on experimental. We have added to this body of information by reporting benchmarking results from a single cell *E. coli* dataset, but more research is needed so that we can get a clearer picture of how well network inference from transcriptomic data works under different conditions, with different organisms and cell types, and using different experimental protocols.

## V. ACKNOWLEDGMENTS

AS acknowledges support from NIGMS-NIH via grant R35GM148351 and from NSF via grant ECCS-1711548.

## REFERENCES

[1] Uri Alon. An introduction to systems biology: Design principles of biological circuits. Chapman and Hall/CRC press, 2020.

[2] M. Bansal, G. D. Gatta, and D. di Bernardo. Inference of gene regulatory networks and compound mode of action from time course gene expression profiles. Bioinformatics, 22(7):815–822, 2006.

[3] Shohag Barman and Yung-Keun Kwon. A Boolean network inference from time-series gene expression data using a genetic algorithm. Bioinformatics, 34(17):i927–i933, 2018.

[4] Attila Becskei and Luis Serrano. Engineering stability in gene networks by autoregulation. Nature, 405(6786):590–593, 2000.

[5] Célia Biane, Franck Delaplace, and Tarek Melliti. Abductive network action inference for targeted therapy discovery. Electronic Notes in Theoretical Computer Science, 335:3–25, 2018.

[6] Sydney B. Blattman, Wenyan Jiang, Panos Oikonomou, and Saeed Tavazoie. Prokaryotic single-cell RNA sequencing by in situ combinatorial indexing. 2019.

[7] A. J. Butte and I. S. Kohane. Mutual information relevance networks: Functional genomic clustering using pairwise entropy measurements. Biocomputing 2000, 1999.

[8] Thalia E. Chan, Michael P.H. Stumpf, and Ann C. Babtie. Gene regulatory network inference from single-cell data using multivariate information measures. Cell Systems, 5(3), 2017.

[9] Francis H. Crick. On protein synthesis. Symposia of the Society for Experimental Biology, 12:138–63, 1958.

[10] Miao Cui, Chao Cheng, and Lanjing Zhang. High-throughput proteomics: A methodological mini-review. Laboratory Investigation, 102(11):1170–1181, 2022.

[11] Ricardo de Matos Simoes, Matthias Dehmer, and Frank Emmert-Streib. B-cell lymphoma gene regulatory networks: Biological consistency among inference methods. Frontiers in Genetics, 4, 2013.

[12] Frank Emmert-Streib, Ricardo de Matos Simoes, Galina Glazko, Simon McDade, Benjamin Haibe-Kains, Andreas Holzinger, Matthias Dehmer, and Frederick Charles Campbell. Functional and genetic analysis of the colon cancer network. BMC Bioinformatics, 15(S6), 2014.

[13] Frank Emmert-Streib, Ricardo de Matos Simoes, Paul Mullan, Benjamin Haibe-Kains, and Matthias Dehmer. The gene regulatory network for breast cancer: Integrated regulatory landscape of cancer hallmarks. Frontiers in Genetics, 5, 2014.

[14] Jeremiah J Faith, Boris Hayete, Joshua T Thaden, Ilaria Mogno, Jamey Wierzbowski, Guillaume Cottarel, Simon Kasif, James J Collins, and Timothy S Gardner. Large-scale mapping and validation of Escherichia coli transcriptional regulation from a compendium of expression profiles. PLoS Biology, 5(1), 2007.

[15] Xin Fang, Anand Sastry, Nathan Mih, Donghyuk Kim, Justin Tan, James T. Yurkovich, Colton J. Lloyd, Ye Gao, Laurence Yang, Bernhard O. Palsson, and et al. Global transcriptional regulatory network for Escherichia coli robustly connects gene expression to transcription factor activities. Proceedings of the National Academy of Sciences, 114(38):10286–10291, 2017.

[16] Nir Friedman, Michal Linial, Iftach Nachman, and Dana Pe’er. Using Bayesian networks to analyze expression data. Journal of Computational Biology, 7(3-4):601–620, 2000.

[17] Socorro Gama-Castro, Heladia Salgado, Alberto Santos-Zavaleta, Daniela Ledezma-Tejeida, Luis Muñiz-Rascado, Jair Santiago García-Sotelo, Kevin Alquicira-Hernández, Irma Martínez-Flores, Lucia Pannier, Jaime Abraham Castro-Mondragón, and et al. Regulondb version 9.0: High-level integration of gene regulation, coexpression, motif clustering and beyond. Nucleic Acids Research, 44(D1), 2015.

[18] Daniel T Gillespie. A general method for numerically simulating the stochastic time evolution of coupled chemical reactions. Journal of Computational Physics, 22(4):403–434, 1976.

[19] Anne-Claire Haury, Fantine Mordelet, Paola Vera-Licona, and Jean-Philippe Vert. TIGRESS: Trustful inference of gene regulation using stability selection. BMC Systems Biology, 6(1), 2012.

[20] Vân Anh Huynh-Thu, Alexandre Irrthum, Louis Wehenkel, and Pierre Geurts. Inferring regulatory networks from expression data using tree-based methods. PLoS ONE, 5(9), 2010.

[21] Vân Anh Huynh-Thu and Guido Sanguinetti. Combining tree-based and dynamical systems for the inference of gene regulatory networks. Bioinformatics, 31(10):1614–1622, 2015.

[22] Vân Anh Huynh-Thu and Guido Sanguinetti. Gene regulatory network inference: An introductory survey. Methods in Molecular Biology, page 1–23, 2018.

[23] P. K. Kreeger and D. A. Lauffenburger. Cancer systems biology: A network modeling perspective. Carcinogenesis, 31(1):2–8, 2009.

[24] Ashwinikumar Kulkarni, Ashley G. Anderson, Devin P. Merullo, and Genevieve Konopka. Beyond bulk: A review of single cell transcriptomics methodologies and applications. Current Opinion in Biotechnology, 58:129–136, 2019.

[25] Tarun Mahajan, Michael Saint-Antoine, Roy D. Dar, and Abhyudai Singh. Limits on inferring gene regulatory networks from single-cell measurements of unstable mRNA levels. 2022 IEEE 61st Conference on Decision and Control (CDC), 2022.

[26] Daniel Marbach, James C Costello, Robert Küffner, Nicole M Vega, Robert J Prill, Diogo M Camacho, Kyle R Allison, The DREAM5 Consortium, Manolis Kellis, James J Collins, and et al. Wisdom of crowds for robust gene network inference. Nature Methods, 9(8):796–804, 2012.

[27] Adam A Margolin, Ilya Nemenman, Katia Basso, Chris Wiggins, Gustavo Stolovitzky, Riccardo Dalla Favera, and Andrea Califano. ARACNE: An algorithm for the reconstruction of gene regulatory networks in a mammalian cellular context. BMC Bioinformatics, 7(S1), 2006.

[28] Patrick E. Meyer, Kevin Kontos, Frederic Lafitte, and Gianluca Bontempi. Information-theoretic inference of large transcriptional regulatory networks. EURASIP Journal on Bioinformatics and Systems Biology, 2007:1–9, 2007.

[29] Thomas G Minchington, Sam Griffiths-Jones, and Nancy Papalopulu. Dynamical gene regulatory networks are tuned by transcriptional autoregulation with microRNA feedback. 2020.

[30] Daniel Moore, Ricardo de Matos Simoes, Matthias Dehmer, and Frank Emmert-Streib. Prostate cancer gene regulatory network inferred from RNA-Seq data. Current Genomics, 20(1):38–48, 2019.

[31] Michael Saint-Antoine and Abhyudai Singh. Limits on inferring gene regulatory networks subjected to different noise mechanisms. bioRxiv, 2023.

[32] Michael M. Saint-Antoine and Abhyudai Singh. Evaluating pruning methods in gene network inference. 2019 IEEE Conference on Computational Intelligence in Bioinformatics and Computational Biology (CIBCB), 2019.

[33] Michael M Saint-Antoine and Abhyudai Singh. Network inference in systems biology: Recent developments, challenges, and applications. Current Opinion in Biotechnology, 63:89–98, 2020.

[34] Lea Schuh, Michael Saint-Antoine, Eric M. Sanford, Benjamin L. Emert, Abhyudai Singh, Carsten Marr, Arjun Raj, and Yogesh Goyal. Gene networks with transcriptional bursting recapitulate rare transient coordinated high expression states in cancer. Cell Systems, 10(4), 2020.

[35] Sydney M. Shaffer, Margaret C. Dunagin, Stefan R. Torborg, Eduardo A. Torre, Benjamin Emert, Clemens Krepler, Marilda Beqiri, Katrin Sproesser, Patricia A. Brafford, Min Xiao, and et al. Rare cell variability and drug-induced reprogramming as a mode of cancer drug resistance. Nature, 546(7658):431–435, 2017.

[36] Claude E. Shannon. A mathematical theory of communication. Bell System Technical Journal, 27(4):623–656, 1948.

[37] Nitin Singh, Mehmet Eren Ahsen, Niharika Challapalli, Hyun-Seok Kim, Michael A. White, and Mathukumalli Vidyasagar. Inferring genome-wide interaction networks using the phi-mixing coefficient, and applications to lung and breast cancer. IEEE Transactions on Molecular, Biological and Multi-Scale Communications, 4(3):123–139, 2018.

[38] Nitin Singh and Mathukumalli Vidyasagar. bLARS: An algorithm to infer gene regulatory networks. IEEE/ACM Transactions on Computational Biology and Bioinformatics, 13(2):301–314, 2016.

[39] Zhong Wang, Mark Gerstein, and Michael Snyder. RNA-seq: A revolutionary tool for transcriptomics. Nature Reviews Genetics, 10(1):57–63, 2009.

[40] Ting Xu, L. Ou-Yang, Xiaohua Hu, and Xiao-Fei Zhang. Identifying gene network rewiring by integrating gene expression and gene network data. IEEE/ACM Transactions on Computational Biology and Bioinformatics, 15(6):2079–2085, 2018.

[41] Bin Zhang and Steve Horvath. A general framework for weighted gene co-expression network analysis. Statistical Applications in Genetics and Molecular Biology, 4(1), 2005.

[42] Haitao Zhao and Zhong-Hui Duan. Cancer genetic network inference using gaussian graphical models. Bioinformatics and Biology Insights, 13, 2019.

[43] Xiaowei Zhu, Mark Gerstein, and Michael Snyder. Getting connected: Analysis and principles of biological networks. Genes amp; Development, 21(9):1010–1024, 2007.

